# High-quality genome assemblies provide clues on the evolutionary advantage of blue peafowl over green peafowl

**DOI:** 10.1101/2023.02.18.529039

**Authors:** Abhisek Chakraborty, Samuel Mondal, Shruti Mahajan, Vineet K. Sharma

## Abstract

An intriguing example of differential adaptability is the case of two Asian peafowl species, *Pavo cristatus* (blue peafowl) and *Pavo muticus* (green peafowl), where the former has a “Least Concern” conservation status and the latter is an “Endangered” species. To understand the genetic basis of this differential adaptability of the two peafowl species, a comparative analysis of these species is much needed to gain the genomic and evolutionary insights. Thus, we constructed a high-quality genome assembly of blue peafowl with an N50 value of 84.81 Mb (pseudochromosome-level assembly), and a high-confidence coding gene set to perform the genomic and evolutionary analyses of blue and green peafowls with 49 other avian species. The analyses revealed adaptive evolution of genes related to neuronal development, immunity, and skeletal muscle development in these peafowl species. Major genes related to axon guidance showed adaptive evolution in peafowl species. However, blue peafowl showed the presence of 42% more coding genes compared to the green peafowl along with a higher number of unique gene clusters, segmental duplicated genes and expanded gene families, and comparatively higher evolution in neuronal and developmental pathways. Blue peafowl also showed longer branch length compared to green peafowl in the species phylogenetic tree. These genomic insights obtained from the high-quality genome assembly of *P. cristatus* constructed in this study provide new clues on the superior adaptability of the blue peafowl over green peafowl despite having a recent species divergence time.

## 1. INTRODUCTION

*Pavo cristatus* is colloquially referred to as peacock or Indian peafowl, and is known for its unique ornamental phenotypes, sexual behaviours, and evolutionary significance. The cultural importance of peacock is not only limited to the Indian subcontinent but expands to Persian, Mesopotamian and ancient Greek cultures. The Asian peafowls belong to the order Galliformes, and family Phasianidae that also includes species like chicken, turkey, quail, etc. Phasianidae family consists of terrestrial, short-winged birds, and has more than 180 species. Indian or blue peafowl had origin in the Indian subcontinent, and shows sexual dimorphism, polygamy, and intricate male display during courtship [1]. Indian peafowl and their closest relative green peafowl, which is the only other species from *Pavo* genus, had diverged around 3 million years ago (mya) [2].

Peacocks have intrigued biologists for hundreds of years and are still a fascinating specimen of study [3,4]. From Charles Darwin’s explanation for the colourful plumage indicating that the vibrant plumage was selected sexually, to Amotz Zahavi proposing his handicap theory [1], a lot has been added to the understanding of peacock’s unique pattern of evolution, still it remains among the most intriguing birds. In addition, factors such as the number of ocelli (eye-spots) in tail [5], certain behavioural factors and calls of peacock had led to the evolution of the distinct traits responsible for sexual selection in peacocks [6]. Gazing pattern of peacocks towards specific display regions plays an important role in intra and inter-sexual selection [7], and in addition the co-evolution of opsin genes and plumage colouration genes is also associated to sexual selection in birds [8]. To perform complex cognitive activities such as sexual selection, brain size has been evolved for better motor control abilities, well developed neural networks, and highly evolved brain organization [9,10]. Comparative genomic analysis also supported sexual selection in peacock, and showed that feather and immune-related gene-pairs underwent selection pressure in this species [4], consistent with Hamilton-Zuk hypothesis [11].

The native habitat of green peafowl was spread across South East Asia, however this species is now extinct or near extinct in Malaysia, Bangladesh, and India, and is also facing reduction in population size in Thailand, Laos, China, and Indonesia [12,13]. Genomic, anthropogenic, and climatic evidences also suggest the decrease in effective population size or endangerment of green peafowl species, and the role of human disturbance in it [3,14]. Habitat loss has confined this species to restricted geographical regions, which caused a reduction in gene flow and higher rate of inbreeding [14].

Blue peafowl has been categorized as species of “Least Concern” whereas green peafowl has been declared as “Endangered” by IUCN for its gradual decrease in population size [15]. The first genome sequencing of blue peafowl (1.16 Gbp) performed by Jaiswal et al. (2018) [3] showed adaptive evolution of genes related to immunity, skeletal muscle, and feather development that aids in phenotypic evolution of blue peafowl, followed by the genome sequencing of this species by other groups [16,17]. The recent genome sequencing of green peafowl revealed a genome size of 1.05 Gbp consisting of 27 pseudochromosomes [14,18]. In this study, we constructed a genome assembly of blue peafowl with the best assembly contiguity till date by performing 10x Genomics sequencing, Oxford Nanopore sequencing, and Illumina short read sequencing, and using previously available Illumina and Nanopore sequencing data of this species [3,16]. Usage of multiple sequencing technologies and hybrid assembly approaches provided better genomic contiguity and helped in better quality of gene space representation, genome annotation, genomic characterization and in revealing novel evolutionary insights to understand the genomic basis of highly valued traits [19–21].

Further, to investigate the genomic basis of their differential adaptability we performed comparative evolutionary analyses of blue and green peafowl species using the high-quality genome assembly of blue peafowl constructed in this study, which revealed adaptive evolution of genes related to neuronal development along with immunity, and skeletal muscle development related pathways in both the peafowl species. However, the genomic evidence highlights better adaptive evolution of blue peafowl species for survival compared to green peafowl.

## 2. MATERIALS AND METHODS

### 2.1 Genome sequencing

The DNA was extracted from the collected blood sample using DNeasy Blood and Tissue kit (Qiagen, CA, USA). DNA was used to prepare a short reads library using NEBNext Ultra II DNA library preparation kit for Illumina (New England Biolabs, England) and sequenced on Illumina HiSeq X instrument (Illumina Inc., USA) for 150 bp paired-end reads. The DNA was amplified using Genomiphi V2 DNA amplification kit. For 10x Genomics linked read sequencing, the amplified DNA library was prepared on a Chromium instrument using Gel Bead Kit v2 and Chromium Genome Library kit (10x Genomics, CA, USA). The prepared library was sequenced on Illumina NovaSeq 6000 instrument (Illumina Inc., USA) for paired-end sequencing. Nanopore long read sequencing library was prepared using the SQK-LSK108 library preparation kit and following the protocol ligation sequencing gDNA. The prepared library was sequenced on a Nanopore sequencer to generate long read data (Oxford Nanopore Technologies, UK).

### 2.2 Genome assembly

#### 2.2.1 10x data assembly

Barcoded 10X Genomics data generated from this study was used for *de novo* genome assembly using Supernova v2.1.1 [22] with default parameters and maxreads=all option without any prior preprocessing, and the haplotype-phased genome assembly was obtained using Supernova mkoutput “pseudohap2” style.

#### 2.2.2 Illumina data assembly

Illumina short read paired-end sequencing data available from the previous study [3] (generated from the same individual), and data generated from this study were used for *de novo* genome assembly. 10x Genomics data from this study was also filtered for barcode sequences using python scripts available in proc10xG (https://github.com/ucdavis-bioinformatics/proc10xG), and used in this assembly. Prior to genome assembly, all three sets of data were quality-filtered using NGSQCToolkit v2.3 [23] with the same parameters used in a previous study [3]. Quality-filtered data were *de novo* assembled using SPAdes v3.15.3 [24] with k-mer value of 101 as used in the previous study [3].

#### 2.2.3 Nanopore data assembly

Oxford Nanopore data generated in this study and previous study [16] were used for adapter trimming using Porechop v0.2.4 (Oxford Nanopore technologies), and the pre-processed reads were used for *de novo* assembly using Flye v2.9 [25]. The assembled genome obtained from Nanopore data was polished three times using Pilon v1.23 [26] with the quality-filtered lllumina short read and 10x Genomics data (barcode-filtered) that were used in the *de novo* assembly performed using SPAdes.

#### 2.2.4 Assembly post-processing and generation of final genome assembly

Three different genome assemblies obtained using three types of sequencing data were scaffolded using the quality-filtered Illumina short read paired-end data from this study and previous studies [3,16], quality-filtered mate-pair data from previous study [16], and the pre-processed Nanopore long read data from this study and previous study [16] using Platanus-allee v2.2.2 [27], separately. The resultant assemblies were further scaffolded with quality-filtered RNA-Seq reads obtained from previous study [28] using AGOUTI v0.3.3 [29]. 10x Genomics reads were used for barcode-processing using Longranger basic v2.2.2 (https://support.10xgenomics.com/genomeexome/software/pipelines/latest/installation), and were used for further scaffolding of the assemblies using ARCS v1.2.2 [30] with default parameters.

The scaffolded assemblies obtained from 10x Genomics data and Illumina short read data were used for assembly merging using Quickmerge v0.3 [31] with 10x Genomics data-based assembly as the hybrid-assembly and minimum alignment length of 4,000 bases. The resultant merged assembly was further scaffolded using the Oxford Nanopore data-based assembly using Quickmerge v0.3 [31] with the previously merged assembly as hybrid-assembly and minimum alignment length of 4,000 bases. The final merged and scaffolded assembly was gap-closed using Sealer v2.1.5 [32] (with k-mer values from 30 to 120 with 10 bp interval), and LR_Gapcloser (three iterations) [33] with the quality-filtered short reads, and the pre-processed Nanopore long reads, respectively, that were used for the *de novo* assembly in the previous steps.

The gap-closed assembly was polished to fix any mis-assembly, erroneous base, or small indel using Pilon v1.23 (three iterations) [26] with the quality-filtered 10x Genomics data (barcode-filtered), and Illumina paired-end short read data that were used for *de novo* assembly in SPAdes. The same quality-filtered short read data was used to further error-correct the Pilon-polished assembly with Seqbug [34] (default parameters), after constructing the individual nucleotide position matrix for each scaffold using bam-readcount (https://github.com/genome/bam-readcount) with the parameters - minimum base quality 25, minimum mapping quality 25, maximum depth 400. Scaffolds with length of ≥ 5,000 bases were extracted to construct the final genome assembly of blue peafowl. To assess the genome assembly completeness, BUSCO v5.2.2 [35] was used with aves_odb10 single-copy orthologous gene set. Also, barcode-filtered 10x Genomics reads (quality-filtered), and quality-filtered Illumina reads from this study and previous study [3] were separately mapped to the final genome assembly of blue peafowl using BWA-MEM v0.7.17 [36] to calculate the read mapping percentage.

Genomic heterozygosity of blue peafowl was estimated using GenomeScope v2 [37] after constructing the k-mer count histogram using Jellyfish v2.2.10 [38] with the quality-filtered Illumina data obtained from this study and the previous study [3]. Sequence variations in blue peafowl genome assembly was identified by mapping barcode-filtered 10x Genomic data (quality-filtered), and quality-filtered Illumina data generated from the same individual using BWA-MEM v0.7.17 [36], and SAMtools v1.9 [39]. BCFtools v1.14 [40] was used for variant calling, and variant filtering was performed with the parameters - variant quality ≥ 30, sequencing depth ≥ 30, mapping quality ≥ 50.

The scaffold-level genome of blue peafowl was assembled into pseudochromosomes by mapping onto green peafowl chromosome-level assembly [18] via genome-wide synteny alignment using Chromosemble in Satsuma v2 [41,42].

### 2.3 Repeat annotation

For identification of repetitive sequences in the final genome assembly [43,44] of blue peafowl, RepeatModeler v2.0.2a was used to construct a *de novo* repeat library [45]. Chicken-specific repeat sequences available in Repbase library [46] were also extracted, and added to the *de novo* repeat library obtained from RepeatModeler for better prediction of repeat sequences in the improved genome assembly of blue peafowl. This combined repeat library was used for soft-masking of the blue peafowl genome assembly using RepeatMasker v4.1.2 (http://www.repeatmasker.org).

### 2.4 Construction of coding gene set

The repeat-masked genome assembly of blue peafowl was used to predict the coding gene sequences using MAKER v3.01.04 [47] with *ab initio* and evidence alignment approaches. Prior to MAKER genome annotation pipeline, quality-filtered RNA-Seq reads from a previous study [28] were used for *de novo* transcriptome assembly of blue peafowl using Trinity v2.13.2 (default parameters) [48]. The protein sequences of blue peafowl, and its phylogenetically closer species - *Gallus gallus, Chrysolophus pictus, Phasianus colchicus, Meleagris gallopavo, Coturnix japonica, Numida meleagris* (belonging to the same Galliformes order) available in Ensembl genome browser 105 [49] were extracted and used as empirical evidence in MAKER pipeline (first round) along with the *de novo* transcriptome assembly obtained from Trinity. The first round of MAKER annotation result was used for training SNAP v2006-07-28 gene prediction program [50], and the second round of MAKER annotation. This result was further used for training AUGUSTUS v3.2.3 gene prediction program [51] for our species and SNAP v2006-07-28 gene prediction program, and the training results were used for a third round of MAKER annotation [52]. After the third or final round of MAKER annotation, coding genes were filtered based on AED (Annotation Edit Distance) value <0.5, and length of ≥ 150 bases to construct the final high-confidence coding gene set of blue peafowl [53]. Additionally, the high-quality genome assembly of blue peafowl was also used for *de novo* prediction of tRNAs and rRNAs using tRNAscan-SE v2.0.7 [54] and Barrnap v0.9 (https://github.com/tseemann/barrnap), respectively, and for homology-based identification of miRNAs using MirGeneDB v2.1 database [55] using BLASTN (95% query coverage and 95% sequence identity).

### 2.5 Collinearity analysis

Pseudochromosome-level assembly of blue peafowl was used to identify the intra-species, and interspecies (between blue and green peafowl) collinear blocks. Collinear blocks for these peafowl species were identified using intra-species and inter-species All-versus-All BLASTP alignments (e-value 10^−5^) using green peafowl protein sequences obtained from a previous study [18], and protein sequences of blue peafowl obtained from this study. MCScanX [56] was used to perform this analysis with the parameter of five genes required to identify a collinear block [57]. Collinear blocks between the longest chromosomes (≥ 5 Mb) of green peafowl and the longest pseudochromosomes (≥ 5 Mb) of blue peafowl were visualized using Evol2Circos [58].

### 2.6 Phylogenetic analysis

Protein sequences of blue peafowl obtained in this study, green peafowl obtained from a previous study [18], and 49 other avian species available on Ensembl genome browser 105 were used to determine the phylogenetic position of the peafowl species across 12 phylogenetic orders.

Protein sequences of the selected species were used to construct the orthogroups using OrthoFinder v2.5.4 [59]. Among these, the fuzzy one-to-one orthogroups containing sequences from all 51 species were identified using KinFin v1.0 [60], and filtered to contain only the longest sequence per species. These filtered orthogroups were individually aligned using MAFFT v7.490 [61], which were concatenated after filtering the empty sites using BeforePhylo v0.9.0 (https://github.com/qiyunzhu/BeforePhylo). The concatenated alignment was used to construct the maximum likelihood dependent species phylogenetic tree using RAxML v8.2.12 [62] with ‘PROTGAMMAAUTO’ substitution model and 100 bootstrap values.

### 2.7 Identification of genes with evolutionary signatures

For the analysis of signatures of adaptive evolution in blue and green peafowl, coding gene information of the six species from Galliformes order available in Ensembl Release 105, green peafowl from previous study [18], and blue peafowl obtained in this study were used. Only the phylogenetically closer species (from Galliformes order itself) were selected for this analysis to identify the more specific genes evolved in this species compared to its closer relatives. Protein sequences of these eight species were used to construct the orthogroups using OrthoFinder v2.5.4 [59], and the orthogroups containing sequences from all eight species were further filtered for the presence of the longest sequence per species. The resultant orthogroups were aligned individually using MAFFT v7.490 [61].

These multiple sequence alignments were used to extract the *Pavo* species-specific genes showing unique amino acid positions compared to the other selected species. In this analysis, any gap and ten positions around the gap present in the alignments were ignored. Functional impact of these substitutions on the protein function was analyzed using Sorting Intolerant From Tolerant (SIFT) [63].

Protein sequence alignments of these orthogroups were used to individually construct maximum likelihood-based gene phylogenetic tree using RAxML v8.2.12 [62] with 100 bootstrap values and ‘PROTGAMMAAUTO’ substitution model. Blue peafowl and green peafowl genes with higher nucleotide divergence compared to genes from other selected species were identified by calculating the branch-length distance values using “adephylo” package available in R [64].

For positive selection analysis, the coding gene sequences (nucleotide) of all eight species were used for orthogroups construction, orthogroups filtering based on the longest sequence per species, and nucleotide sequence alignment using MAFFT v7.490 [61]. The nucleotide alignments were converted into PHYLIP format, and used for positive selection analysis using “codeml” program from PAML v4.9a [65], along with the species phylogenetic tree across the eight species (constructed in a similar manner as described earlier). Likelihood-ratio tests were carried out, and blue peafowl and green peafowl genes qualifying against null model with FDR-corrected p-values <0.05 in chi-square analysis were extracted as genes showing positive selection. Further, genes from both the peafowl species containing codon sites with >95% probability for the foreground lineage obtained from Bayes Empirical Bayes (BEB) analysis were identified as genes with positively selected codon sites.

Genes showing at least two of the three evolutionary signatures – higher nucleotide divergence, positive selection, and unique substitution with functional impact were termed as genes with Multiple Signs of Adaptive evolution (MSA) [66,67].

### 2.8 Evolution of gene families

Evolution of gene families in blue and green peafowl species in terms of gene family expansion or contraction was analyzed using CAFÉ v4.2.1 [68] with respect to 49 other avian species used for species phylogenetic tree construction in this study. All-versus-All BLASTP homology search result using the longest isoforms of the protein sequences of these species was clustered using MCL v14.137 [69], and gene families filtering was performed as suggested for CAFÉ analysis. The resultant gene families along with the ultrametric species phylogenetic tree that was obtained using the divergence time between blue and green peafowl (from TimeTree database) were used for CAFÉ analysis with two-lambda (λ) model, where species from Galliformes order were assigned a separate λ-value from the other selected species.

### 2.9 Exon expansion analysis

Exon expansion in the coding gene sequences of the two peafowl species with respect to each other species was analyzed. For identification of the orthologous genes in these two species, the protein sequences of these two species were used to construct the orthogroups using OrthoFinder v2.5.4 [59]. Only the longest protein sequence per species was retained in each orthogroup. Number of exons expanded or contracted in the orthologous genes of one peafowl species compared to the other was calculated from the respective GFF files.

### 2.10 Functional annotation

The coding gene set of blue peafowl was mapped against NCBI-nr database using BLASTP with e-value cutoff 10-5, and the unmapped genes were mapped against Pfam-A [70] and SwissProt [71] databases. Protein function prediction for genes without any match against all three databases was performed using InterProScan v5.54-87.0 [72]. Genes that showed evolutionary signatures in both the peafowl species were annotated against KAAS v2.1 web server [73] and eggNOG-mapper v2 [74]. Shared orthologous clusters or gene families among blue peafowl and its phylogenetically closer species (according to the species phylogenetic tree) from Galliformes order were identified using OrthoVenn2 [75].

## 3. RESULTS

### 3.1 Genome assembly

A total of 502 Gb (444.3X) of Illumina short read and mate-pair data, 91.8 Gb (81.2X raw sequencing coverage) of 10x Genomics linked read data, and 8.6 Gb (7.6X) of Oxford Nanopore data was used to construct a high-quality genome assembly of blue peafowl. A total of 92.7 Gb of transcriptome data of this species from a previous study [28] was also used in downstream analysis to help in the genome assembly and gene prediction. The separate assemblies constructed using 10x Genomics, Illumina short reads, and Nanopore data were used to build the final scaffolded and merged genome assembly of blue peafowl (≥ 5 Kbp) with a genome size of 1.13 Gbp (comprised of 1,665 scaffolds), N50 value of 4.95 Mb, longest scaffold size of 17.6 Mb. After the single scaffolded assembly was constructed using Quickmerge, base correction was performed in the Pilon-polished assembly using SeqBug [34], which corrected 0.002% base positions. Further, 89.5% BUSCOs could be found in the final genome assembly. 93.75% barcode-filtered 10x Genomics linked reads from this study, and 93.82% quality-filtered Illumina short reads from this study and a previous study [3] were mapped on the assembled genome, attesting to the good quality of the genome assembly. The blue peafowl genome was estimated to contain 0.47% heterozygosity. Sequence variation analysis in the final assembled genome showed that 0.17% of the base positions had single nucleotide variations that were present across 1,471 scaffolds.

This genome assembly was further improved by constructing a pseudochromosome-level assembly. 438 superscaffolds and 69 pseudochromosomes were constructed from the scaffolds of blue peafowl genome assembly in this study, and this pseudochromosome-level assembly comprised of 1.13 Gbp with an N50 value of 84.81 Mb and covering 89.1% BUSCOs. The assembled genome size was same as the previously estimated genome size of 1.13 Gbp [3]. The pseudochromosome-level genome assembly statistics were significantly improved than the previously available *P. cristatus* genome assemblies [3,16,17] (**Table 1**).

**Table 1.**
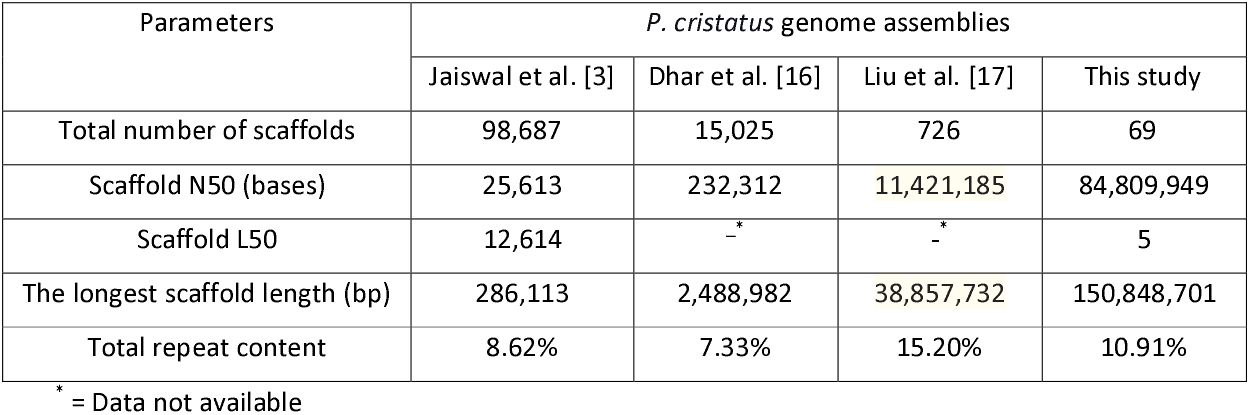
*P. cristatus* pseudochromosome-level assembly statistics comparison with previous studies

### 3.2 Genome annotation

Repeat-masking of the blue peafowl genome assembly was performed using a combined repeat library comprising of 206 chicken-specific repeat sequences and 348 *de novo* constructed repeat family sequences obtained from RepeatModeler. 9.87% of the blue peafowl pseudochromosome-level genome assembly consisted of interspersed repeats (5.51% L2/CR1/Rex type of retroelements, 0.86% DNA transposons) and 0.82% of the genome was found to contain simple repeats. LINEs were the most prevalent repeat elements in blue peafowl genome, similar to green peafowl [18].

Prior to coding gene prediction, *de novo* transcriptome assembly using previously available data identified 904,608 assembled transcripts (N50 value = 3,873 bp), that were used as empirical evidence in MAKER pipeline. Coding genes prediction using the repeat-masked genome assembly with MAKER genome annotation pipeline identified a total of 25,681 coding gene sequences after three rounds of comprehensive MAKER annotation, and AED value cut-off and length-based filtering. In contrast, the recently sequenced genome of green peafowl contained only 14,935 genes with a higher percentage of genomic repeats (15.92%) [18]. 83.3% genes of the final blue peafowl coding gene set could be annotated using the publicly available databases and InterProScan, which is higher than the previously reported study [17]. 290 tRNAs, 239 miRNAs, and 28 rRNAs were also identified in blue peafowl genome assembly.

### 3.3 Collinearity and orthologous gene clustering

Intra-species and inter-species collinearity analysis using the genome assemblies of blue and green peafowls revealed 22.44% of blue peafowl and 8.09% of green peafowl coding genes to be involved in the intra-species collinear blocks of the two respective species. 583 inter-species collinear blocks between the two peafowl species were also identified, which included 48.88% and 74.52% of the coding genes of blue and green peafowls, respectively. The absence of a major fraction of blue peafowl coding genes in the inter-species collinear blocks point towards a higher number of unique coding genes in this species. The inter-species collinear blocks between the longest chromosomes of the two *Pavo* species (≥ 5 Mb) are visualized in **Figure 1**.

**Figure 1.**
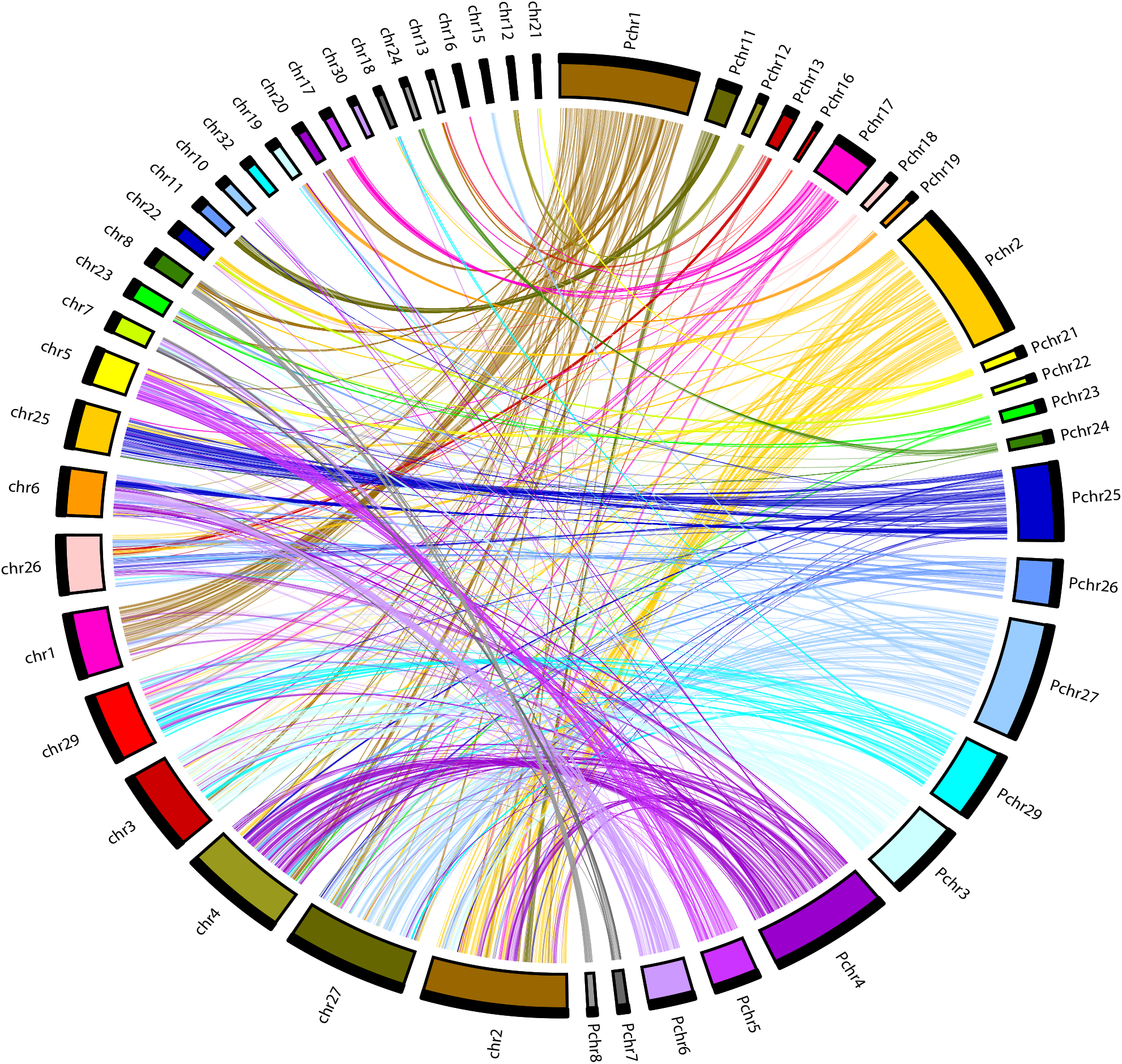
Circular plot showing collinear blocks identified between the chromosomes of green peafowl and the pseudochromosomes of blue peafowl. Right side of the circle represents the pseudochromosomes of blue peafowl (Pchr = Pseudochromosome), and left side of the circle represents the chromosomes of green peafowl (chr = chromosome).

Further, the number of gene clusters were found to be similar (13,210 - 14,301) in blue peafowl, green peafowl, and other species from Galliformes order, however a higher number of unique clusters were observed in blue peafowl species (**Figures 2A-2B**) that also supports the presence of a higher number of unique coding genes in blue peafowl.

**Figure 2.**
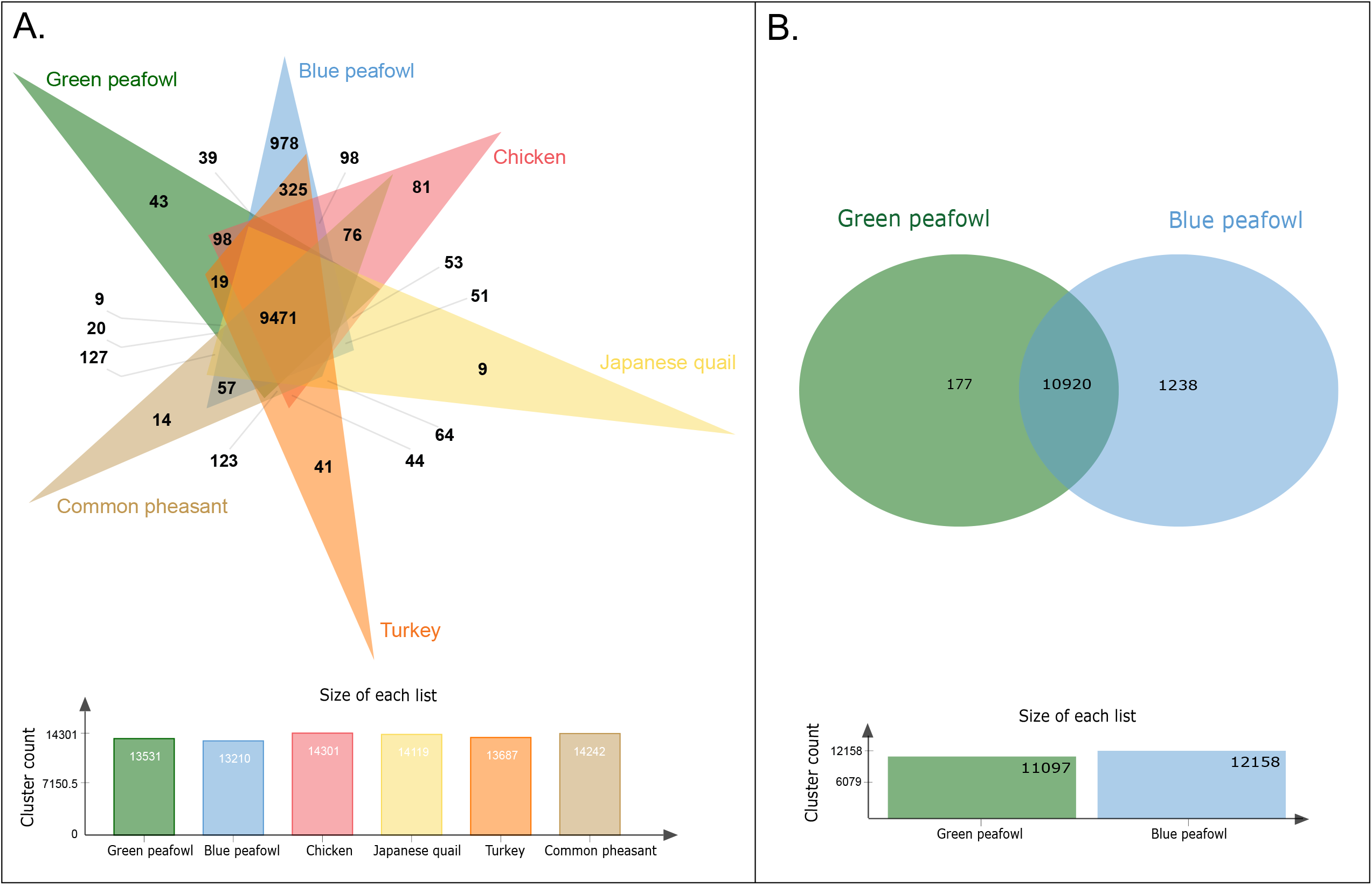
Shared orthologous gene clusters obtained from OrthoVenn2 analysis. **A**. Shared orthologous clusters among blue peafowl and five other phylogenetically closer birds from Galliformes order, **B**. Shared orthologous clusters between blue and green peafowl species.

### 3.4 Phylogenetic position

Phylogenetic position of the two peafowl species was determined with respect to 49 other avian species available in Ensembl genome browser 105 [49]. A total of 1,441 fuzzy one-to-one orthogroups were identified across all 51 selected bird species, and 899,247 sequence alignment positions were used to construct the species phylogenetic tree. Blue and green peafowl belonging to the *Pavo* genus were positioned in the same clade and were found to be the closest to each other in the phylogenetic tree (**Figure 3**). However, blue peafowl had longer branch length than green peafowl in the species phylogenetic tree. Among other species from Galliformes order, *Gallus gallus* was the closest phylogenetically placed species of these peafowl species, and *Numida meleagris* had diverged the earliest. Species from Anseriformes order were phylogenetically closest to the species from Galliformes order compared to species from other avian phylogenetic orders.

**Figure 3.**
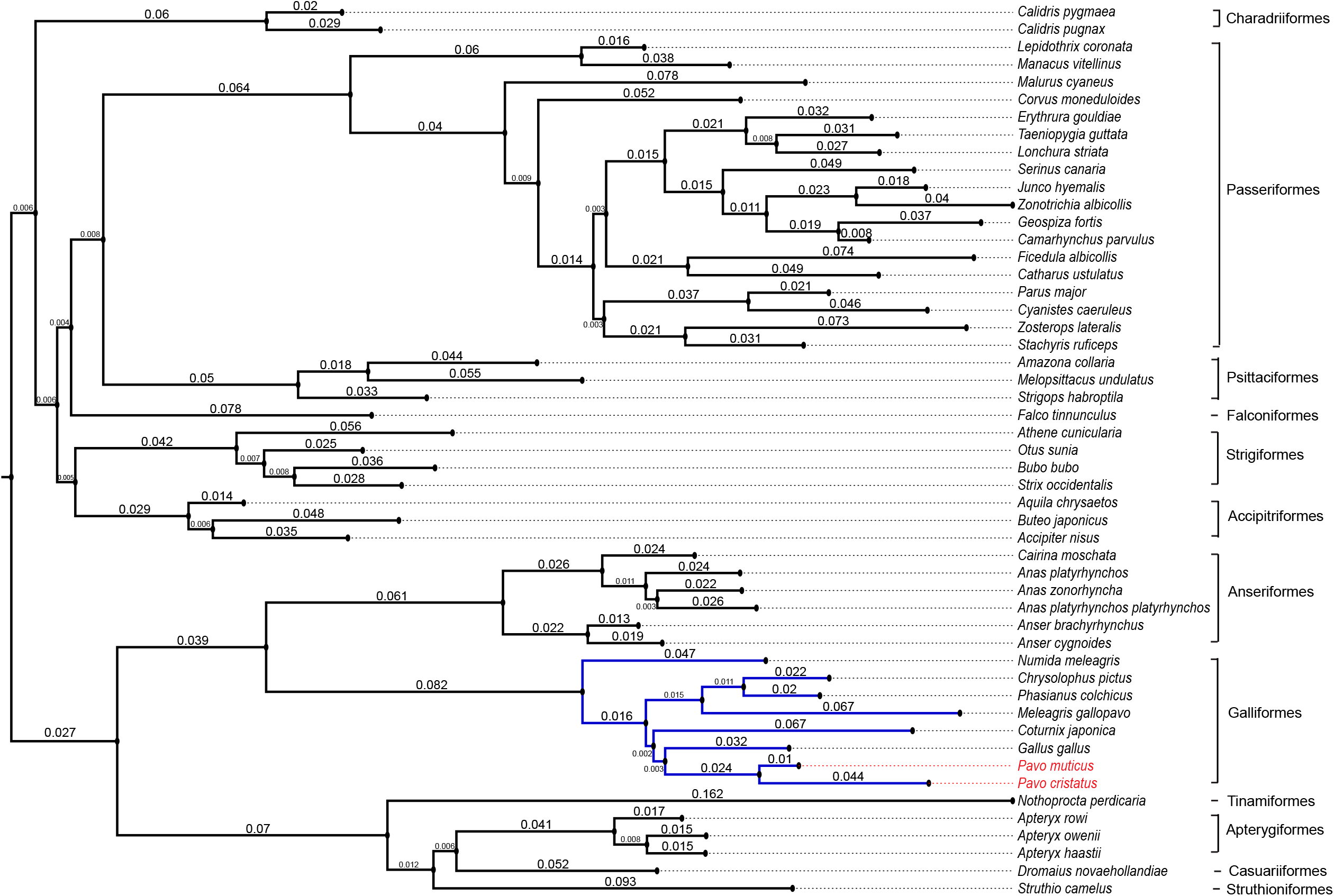
Phylogenetic position of the peafowl species with respect to 49 other avian species available in Ensembl release 105. The numbers mentioned in the phylogenetic tree represent the branch length values.

### 3.5 Genome-wide exon expansion

The analysis of orthogroups (containing genes of the two peafowl species) revealed 4,119 genes in blue peafowl, and 3,960 genes in green peafowl that showed expansion in exon numbers with respect to each other. 16 genes in each of the two *Pavo* species showed ≥ 10 times expansion in terms of exon number along with increased coding gene length and decreased average exon length in most cases. Among these genes, titin had the highest number of expanded exons in blue peafowl (117 exons in blue peafowl and 4 exons in green peafowl). Titin provides structural continuity of a sarcomere in vertebrate striated muscles, and thus helps in contraction of avian flight muscle [76].

In contrast, *COL11A1* showed the highest number of expanded exons in green peafowl (3 exons in blue peafowl and 65 exons in green peafowl) followed by another collagen gene *COL24A1*. Collagens are responsible for structure and strength of connective tissues to support body parts such as cartilage, muscles, and organs [77].

### 3.6 Evolution of gene families

A total of 12,107 filtered (size and clade-based filtering) gene families were identified across the 51 avian species used for species phylogenetic tree construction. In blue peafowl, 4,382 and 1,895 gene families were expanded and contracted, respectively. In green peafowl, 576 gene families were expanded, and 1,717 families were contracted. 174 and 75 highly expanded gene families (with >10 expanded genes) were obtained in blue and green peafowl, respectively, among which 69 were common in both the peafowl species. Furthermore, these 69 highly expanded common gene families were majorly involved in neuronal and signalling KEGG pathways - axon guidance, pathways of neurodegeneration, cytokine-cytokine receptor interaction, chemokine signalling pathway, Hippo signalling pathway, signalling pathways regulating pluripotency of stem cells, neuroactive ligand-receptor interaction, endocytosis, Rap1 signalling pathway, TGF-beta signalling pathway, Ras signalling pathway, cholinergic synapse, Melanogenesis, MAPK signalling pathway, Wnt signalling pathway, regulation of actin cytoskeleton, Phospholipase D signalling pathway, and others. However, in terms of gene numbers, the highly expanded common gene families in blue peafowl contained a larger number of genes compared to green peafowl.

### 3.7 Genes with evolutionary signatures

For the identification of evolutionary signatures in the two peafowl species, a total of 8,527 orthogroups were constructed across eight species from Galliformes order. Compared to the other species, 1,077 genes showed higher nucleotide divergence, 605 genes were positively selected (p-values <0.05), and 700 genes contained unique amino acid substitutions with functional impact in blue peafowl. Among these genes, 429 genes were identified as MSA genes, and 41 genes showed all three evolutionary signatures. In green peafowl, 174 genes showed higher nucleotide divergence, 660 genes showed unique substitution with functional impact, and 142 genes showed positive selection (p-values <0.05), among which 110 genes were MSA genes, and 17 genes displayed all three evolutionary signatures.

The MSA genes of blue peafowl were involved in KEGG pathways including neurodegeneration, endocytosis, axon guidance, protein processing in endoplasmic reticulum, PI3K-Akt signalling, and other pathways. The MSA genes of green peafowl were involved in KEGG pathways - MAPK signalling pathway, Ras signalling pathway, calcium signalling pathway, PI3K-Akt signalling pathway, endocytosis, neuroactive ligand-receptor interaction, and others. Among the genes that showed all three evolutionary signatures, genes related to neuronal development, immune response, and cytoskeletal functions were prominent in both the peafowl species.

#### 3.7.1 Adaptive evolution in nervous system development-related genes in peafowl species

Genes associated with axon guidance, neuronal differentiation, axon regeneration and neurite outgrowth activity, facilitation of sodium-activated potassium channel activity in neurons, and neuropeptide signaling showed all three evolutionary signatures in *P. cristatus*. Among the key *P. muticus* genes showing all evolutionary signatures, genes involved in development of neural cells, synapse and dendrites, axon guidance, myelination and development of Schwann cells were notable.

Among the highly expanded gene families that are common in both *Pavo* species, gene families involved in axon guidance, Notch signaling, and other neuronal processes are noteworthy. However, in the highly expanded common gene families, gene families showed more expansion in blue peafowl compared to green peafowl in terms of gene numbers.

#### 3.7.2 Adaptive evolution in immunity and cytoskeletal genes in Asian peafowls

Immunity-related genes were also found among the genes showing all evolutionary signatures and had highly expanded gene families. Among the *P. cristatus* genes with three evolutionary signatures, genes associated with Ras signaling, MAPK signaling, and Toll-like receptor (TLR) signaling, were notable. Among the *P. muticus* genes with three evolutionary signatures, genes involved in functions such as immune cells development were present.

In these *Pavo* species, extracellular matrix genes, and genes related to muscle functioning were found among the genes with all three evolutionary signatures, and highly expanded gene families.

Cytoskeletal activities related gene families such as gap junction protein family, myosin, claudin, dynein heavy chain, calpain, cadherin, and actin were noteworthy among the highly expanded gene families of both *Pavo* species.

#### 3.7.3Adaptive evolution in feather development and visual genes

Among the genes showing evolutionary signatures in blue peafowl, genes involved in feather color determination were present. Further, feather development and melanin deposition-related genes also showed exon expansion in these *Pavo* species. Further, genes from highly expanded gene families common in both *Pavo* species, positively selected genes of both *Pavo* species, and MSA genes of blue peafowl were involved in melanogenesis (KEGG pathway). Additionally, genes associated with TGF-β, Wnt, and MAPK signaling pathways that are also involved in feather development were found among the adaptively evolved genes, in accordance with the previous studies [3,17].

Birds have evolved their visual system as they heavily rely on the same to adapt in various light conditions [8]. Among the opsin genes that developed in a non-neutral way in birds [8], three genes had greater number of exons in blue peafowl compared to green peafowl, and one gene showed exon expansion in green peafowl.

## 4. DISCUSSION

The peafowl species are known for their unique phenotypic characteristics and evolutionary importance. In this study, the high-quality genome assembly of blue peafowl obtained using Illumina short reads, 10x Genomics sequencing, and Oxford Nanopore long read technology helped in constructing a pseudochromosome-level assembly consisting of 69 pseudochromosomes [18] with the highest N50 value (84.81 Mb) of blue peafowl genome till date. This high-quality assembly helped in gaining important evolutionary and comparative insights on the two peafowl species and can be used as a valuable reference genome for future studies of these intriguing bird species.

The final coding gene set of blue peafowl was constructed using a combination of *ab initio* and homology-based approach with AED value and length-based filtering criteria to ensure a good quality gene set. A comprehensive genome-wide phylogenetic tree constructed with all available avian species in Ensembl 105 along with green peafowl species using this high-confidence coding gene set helped in better resolution of phylogenetic position of peafowls and showed that the clade formed by both *Pavo* species was closest to *Gallus gallus* among other Galliformes order species. This observation provides further support to the previous studies [3,16], except a study where *P. cristatus* was closer to *Meleagris gallopavo* perhaps due to the inclusion of lower (15 species) number of species in their phylogenetic tree [17].

The genes related to pathways of immunity and cytoskeleton were found to be adaptively evolved in blue peafowl in previous studies [3,17]. Comparative evolutionary analyses of blue peafowl, green peafowl, and six other species from the Galliformes order in this study showed the genes majorly involved in neuronal development along with immunity, skeletal muscle development, feather, and visual system development to be adaptively evolved in both the peafowl species. Adaptive evolution of immune response-related genes is responsible for the immunocompetence of peafowl species required for sexual selection, whereas skeletal muscle and feather development explain their large body size and decorative feathers [3,17]. However, in this study the key genes related to neuronal development showed evolutionary signatures, gene family expansion and exon expansion that provide additional genomic insights into the adaptive evolution in the two peafowl species.

For performing specialized cognitive activities such as sound localization, foraging, and assessing rival individuals, avian species possess complicated neural networks and increased brain size, which makes birds as one of the brainiest evolved organisms among the vertebrates [10,78]. In addition, males and females evaluate their mates, and males also assess their rival males through gazing that is one of the most important cognitive functions in peacock and examples of intersexual and intrasexual selection activities [7], for which they would require an evolved nervous system. To support this, one of the key results from this study showed adaptive evolution of genes involved in functioning of nervous system such as axon guidance and neuronal differentiation in peafowl species. Neuron functioning largely relies on development of proper axonal connections, which depends on attractive or repulsive forces generated by the binding of guidance molecules with the receptors present on growth cones, a phenomenon known as axon guidance [79]. Extracellular matrix proteins such as cadherin, and extracellular developmental proteins such as Wnt are also involved in axon guidance and synapse formation in the nervous system [80,81], and were highly expanded in both peafowl species. Besides this, adaptively evolved genes related to neuronal differentiation in different regions of brain were also identified in these peafowl species. Taken together, these observations indicate adaptive evolution in nervous system in peafowl species that perhaps helps in sexual selection [4] and other complex cognitive activities.

One of the interesting observations from this study was the increase in exon-intron numbers in the orthologous genes of the two peafowl species that perhaps occurred due to inclusion of new exons which can arise from splice motifs or upstream shortened introns [82], and gain of exons could be associated with increased gene expression by activating new transcription start sites (TSSs) – a phenomenon known as Exon-mediated activation of transcription starts (EMATS) [83]. The exonintron expansion was also prominent in opsin and plumage colouration genes that can be associated with the distinct phenotypic characteristics of peafowl, along with nervous system and immunity-related genes. Previous studies have suggested an adaptive association between opsin and plumage colouration genes, which is linked to the advantages during sexual selection [8]. This corroborates well with the case of peafowl species, given the importance of visual cues in sexual selection in peafowls [7].

The two Asian peafowl species, green peafowl and blue peafowl provide an intriguing example of differential adaptability where the former is left with a limited number of individuals and is an “Endangered” species, whereas the latter is a species of “Least Concern”. This study provides genomic and evolutionary clues for the difference in the survival and existence status of the two peafowl species (**Figure 4**). Although blue and green peafowl had a species divergence time of only 3 mya, the striking contrast in number of coding genes present in blue peafowl (25,681 genes) compared to green peafowl (14,935 genes [18]) is indeed intriguing. This perhaps can be explained by the presence of a higher percentage of segmental duplicated genes, a greater number of expanded gene families, a lesser percentage of genes present in inter-species collinear blocks, and a higher number of unique gene clusters in blue peafowl (**Figure 2B**) compared to green peafowl.

**Figure 4.**
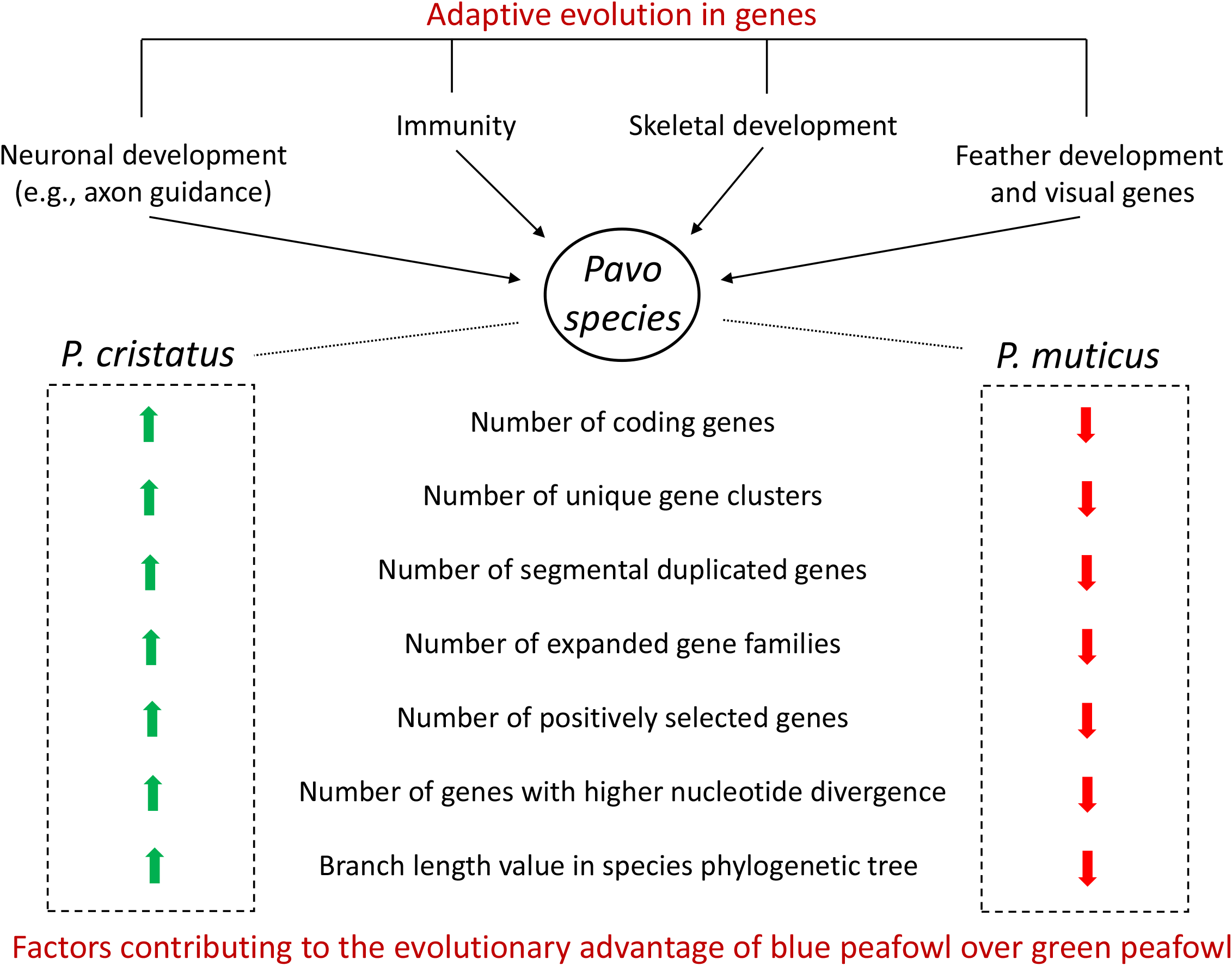
Comparative genomic and evolutionary characteristics of the two peafowl species.

Further, a larger number of genes showed higher nucleotide divergence and positive selection in blue peafowl compared to green peafowl, indicating adaptive evolution to be more prominent in blue peafowl. The signatures of adaptive evolution in genes associated with nervous system development, immunity, and other functions were also higher in blue peafowl. It was also supported by the phylogenetic analysis where the branch length was observed to be comparatively longer in case of blue peafowl compared to green peafowl suggesting a higher rate of evolution in blue peafowl species (**Figure 3**). These findings provide valuable clues on the better adaptability and survival of the blue peafowl compared to the green peafowl. Besides these, PSMC results suggested a more recent second population bottleneck event in green peafowl (~20,000-10,000 years ago) [14] compared to blue peafowl (~450,000 years ago) [3]. The effective population size of green peafowl experienced a steep decline during the recent second population bottleneck event, whereas blue peafowl population was comparatively stable [3,14].

The social behavior and adaptability of the two Asian peafowls also appear to contribute to their contrasting populations sizes. The impact of habitat loss and exploitation by humans for food and commercial purposes seems to have affected the green peafowl population more since it is less tolerant to human activities. A reduction in population contributes to gene flow reduction, high inbreeding, low genetic diversity, and thus leads to a higher extinction possibility [14]. Thus, despite having a recent divergence time, the above-mentioned factors seem to have contributed to the distinct genomic divergence of these peafowl species, and a reduction in population size of green peafowl. This was also the case in Felidae family that evolved in eight lineages over a time period of only six million years [84], where cheetah showed lower genetic diversity and faced a population decline compared to other felid species mostly because of a recent population bottleneck event, loss of habitat, and difficulties in captive breeding, similar to the case of green peafowl [84,85].

Taken together, it is tempting to speculate that the survival and adaptability of an organism appears to be a complex interplay of adaptive evolution, social behavior and environmental influences, and the case of the two peafowls is one such intriguing example that needs more studies to determine the quantitative impacts of these factors. Further, the high-quality genome assembly of *P. cristatus* constructed in this study will act as a valuable reference for future studies of these intriguing bird species.

## 5. CONCLUSION

In this study, we constructed a high-quality assembly of blue peafowl (*P. cristatus*) genome and performed comparative analyses of blue and green peafowl species to understand the genomic basis of differential adaptability of these two species. This study revealed adaptive evolution of nervous system developmental genes along with immunity, and skeletal muscle development genes in both the peafowl species. However, blue peafowl showed better adaptive evolution compared to green peafowl in presence of higher number of expanded gene families, segmental duplicated genes, unique gene clusters, and genes with evolutionary signatures, which highlights the distinct genomic divergence and provides genomic clues on the contrasting population size of the two Asian peafowl species.

## AUTHORS’ CONTRIBUTIONS

VKS conceived and coordinated the project. SM prepared the samples for genome sequencing. AC and VKS designed computational framework of the study. AC and SMD performed genome assembly, genome annotation, synteny analysis, and OrthoVenn analysis. AC performed phylogenetic analysis, CAFÉ analysis, exon expansion analysis, analysis of genes with evolutionary signatures, and performed the functional annotation of gene sets. AC and SMD constructed the figures. AC and VKS interpreted the results. AC, VKS, and SMD wrote the manuscript. All the authors have read and approved the final version of the manuscript.

## ACKNOWLEDGEMENTS

AC and SM thank Council of Scientific and Industrial Research (CSIR) for fellowship. SMD thanks DST-INSPIRE for providing fellowship. The authors thank the NGS facility at IISER Bhopal and the intramural research funds provided by IISER Bhopal. We thank Dr. Atul Gupta, Wildlife Veterinary Officer and Director of Van Vihar National Park, Bhopal for providing the blood samples of peacock. We thank Dr. Tista Joseph and Dr. Niraj Dahe, Wildlife Veterinary Officers at Van Vihar National Park for their help in sample collection for the first peacock genome sequencing project. The authors also thank Dr. Rituja Saxena, Dr. Shubham K. Jaiswal, and Dr. Ankit Gupta for their scientific inputs from the first peacock genome sequencing project.

## ETHICS APPROVAL

This study was approved by the Institute Ethics Committee, IISER Bhopal.

